# Task Difficulty and Limb Dominance Modulate the Effects of Ageing on Neuromuscular Function

**DOI:** 10.1101/2025.09.01.673460

**Authors:** Yuxiao Guo, Eleanor J. Jones, Abdulmajeed Altheyab, Nishadi N. Gamage, Bethan E. Phillips, Philip J. Atherton, Mathew Piasecki

## Abstract

**Background:** Neuromuscular function is critical for independence in ageing, yet asymmetries between dominant and non-dominant limbs, arising from central or peripheral mechanisms, are not well understood. This study examined age- and limb-related differences and motor unit (MU) firing behaviour of the vastus lateralis under tasks of varying difficulty.

**Methods:** Twenty-one young (22 ± 4 years; 15M, 6F) and seventeen older adults (74 ± 5 years; 12M, 5F) performed constant and variable force unilateral isometric knee extensions. In both limbs, high-density surface electromyography signals were decomposed into MU spike trains. Force control and MU firing properties were analysed using multilevel mixed-effects regression models.

**Results:** Older adults showed reduced maximal muscle strength (p<0.001) and increased force tracking error (p=0.008), MU firing rate (MUFR) was significantly lower in older adults during constant contractions (p = 0.001) and trended toward lower during variable contractions (p=0.061). MUFR variability showed a significant Leg × AgeGroup interaction (p<0.001); older adults had greater variability in non-dominant legs, while younger adults showed the opposite. With variable force contractions in both age groups, MUFR was higher during ascending segments with greater variability during descending segments.

**Conclusion:** Neuromuscular ageing involves asymmetric adaptations rather than a uniform decline, with leg dominance effects become more pronounced under variable force modulation. Task difficulty amplifies these asymmetries, underscoring the need to consider limb-specific neural control in age-related motor assessments.

## Introduction

Inter-limb asymmetry refers to imbalances in performance between opposing limbs, arising from bilateral asymmetry of descending motor output between the right and left hemispheres (Kapreli *et al*., 2006). Other factors may also contribute to the development of such imbalances, including biomechanical disparities, previous injuries (Kuenze *et al*., 2015) or disease states (Larson *et al*., 2013), and the effects of ageing (McGrath *et al*., 2021). Consequently, such imbalances can give rise to a favoured limb when performing tasks that require strength or control (Newton *et al*., 2006). This has clear implications in ageing as grip strength asymmetry in older people is a strong predictor of future falls (McGrath *et al*., 2021), and older females who experienced falls exhibited significantly greater bilateral strength differences than those who did not (Skelton *et al*., 2002).

Wider evidence pertaining to limb strength asymmetry is conflicting and varies between muscle groups (Rahnama *et al*., 2005; Lanshammar & Ribom, 2011; Lathrop-Lambach *et al*., 2014), with one study demonstrating significant asymmetry in the knee extensors and flexors, showing ∼13% greater strength during isokinetic contractions in dominant side compared to non-dominant side, and a further showing 9% greater force in the dominant arm flexors of young (Lecce *et al*., 2025). However, a comprehensive review of studies examining the comparative strength of the knee extensors as well as their influence on performance in jump tasks, revealed no significant disparities between the limbs (McGrath *et al*., 2016). The presence of such muscle asymmetry can have implications for the interpretation of clinical evaluations of skeletal muscle and for research studies that employ unilateral intervention models, and explanatory mechanisms are not fully established. Functionally, strength asymmetries in the lower limbs have been associated with impaired mobility, gait instability, and a higher risk of falls (Mertz *et al*., 2019; Si *et al*., 2024), particularly in older adults. Understanding whether such asymmetries arise from neural or peripheral factors is therefore clinically relevant.

The control and coordination of neural inputs play a crucial role in achieving graded increases in muscle force. This involves concurrent variations in the number of active motor units (MU) (known as MU recruitment) and modulation of the discharge frequency of action potentials within these units (known as rate coding) (Del Vecchio *et al*., 2019), probably leading to the uniformly distributed muscle strength and motor control ability (Enoka & Farina, 2021) across limbs. The evidence regarding disparities in MU recruitment and firing related to laterality is also inconclusive. A study investing MU discharge properties in a hand muscle during 30% of maximal voluntary contraction (MVC) revealed dissimilar MU characteristics between limbs with the dominant hand having lower firing rates (FR), FR variability and recruitment threshold when compared with the non-dominant hand (Adam *et al*., 1998). A more recent study of unimanual movements showed less common synaptic inputs to motoneurons serving intrinsic and extrinsic muscles of the dominant hand, interpreted as enabling a higher flexibility in MU recruitment (Maillet *et al*., 2022). Another investigation demonstrated higher MUFR and a greater proportion of common synaptic inputs in dominant biceps (Lecce *et al*., 2025). Yet in lower limbs, there were similar levels of MUFR and MUFR variability bilaterally across submaximal isometric contractions at 10, 20, 40 and 60% of MVC in tibialis anterior (Petrovic *et al*., 2022).

The existing body of knowledge primarily includes young adults, leaving a lack of information regarding the influence of age on limb dominance, particularly for the quadriceps. As the primary extensors of the knee joint, the quadriceps are essential for postural stability, gait propulsion, and fall prevention, making them a key muscle group for examining functional asymmetry in the lower limbs and one of the most susceptible to atrophy with ageing (Maden-Wilkinson *et al*., 2013). Importantly, many studies of ageing MUs have focused only on unilateral movements, and any implications arising from the presence of limb strength asymmetry, as well as central drive and peripheral motor unit properties, as a result of increasing age are yet to be determined.

The purpose of this study was to investigate age- and limb-related differences in the neuromuscular function of the knee extensors, focusing on muscle strength, force control ability, and vastus lateralis MU characteristics across tasks of varying difficulty in young and older adults.

## Materials and Methods

### Ethics and Participants

This research was approved by the University of Nottingham Faculty of Medicine and Health Sciences Research Ethics Committee (FMHS-160-0121, 90-0820, 390-1121) and was conducted between 2022 and 2024 in accordance with the Declaration of Helsinki.

A total of 38 healthy individuals were recruited from the local community through advertisements. Participants were allocated to two age-defined groups based on predetermined criteria: a younger group (target range18-35 years) and an older group (target range 65-85 years).

Prior to enrolment, participants completed a comprehensive clinical screening examination at the University of Nottingham, Royal Derby Hospital site and subsequently provided written informed consent. Participants were excluded if they had BMI< 18.5 or > 35 kg/m^2^; were competitive in any athletic discipline at county level or above; had a musculoskeletal disorder; respiratory disease; neurological disorder; metabolic disease; active cardiovascular problems; active inflammatory bowel or renal disease; recent steroid treatment within 6 months or hormone replacement therapy.

Leg dominance was determined by asking participants which leg they would use to kick a ball, which has been reported to have a strong agreement with observed leg dominance during bilateral mobilizing and unilateral stabilizing tasks (van Melick *et al*., 2017). Participants were instructed to abstain from any vigorous physical exercise for at least 48 hours prior to testing and to avoid caffeine and alcohol consumption for 24 hours before the testing session. All testing sessions were conducted in the morning, with participants arriving at the laboratory at 0900h in a fasted state.

### Experimental protocol

Each participant completed a single experimental session that included the assessment of maximal voluntary isometric contraction (MVC) torque of the knee extensors, a familiarisation task, and the evaluation of muscle force control. Throughout these procedures, high-density surface electromyography (HDsEMG) signals were recorded from the vastus lateralis muscle (Figure 1). All measurements were performed bilaterally.

**Figure 1.**
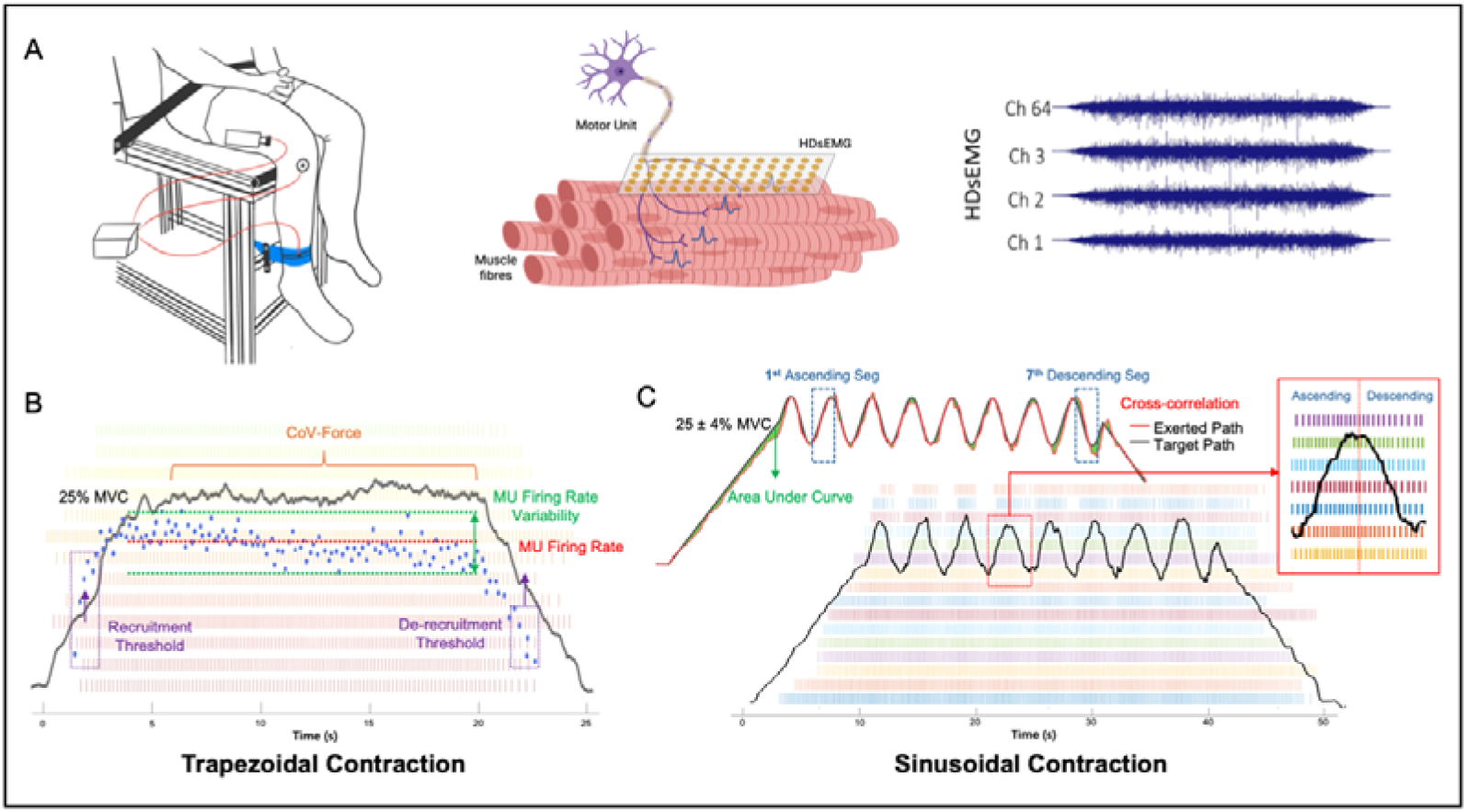
Experimental approach. (A) A participant is seated with the ankle securely attached to a load cell via a non-compliant strap. The load cell measured the force produced by the knee extensors during isometric contractions. High-density surface electromyography (HDsEMG) with a 64-channel grid was placed over the belly of the vastus lateralis. (B) Representative data from the trapezoidal contraction at 25% of maximal voluntary contraction (MVC). Force steadiness (CoV-Force) and motor unit (MU) firing characteristics are shown, including recruitment and de-recruitment thresholds (purple arrow), MU firing rate (red dashed line), and firing rate variability (coefficient of variation of inter-spike interval; green dashed lines). (C) Representative data from the sinusoidal force-tracking task (target force: 25 ± 4% MVC). Tracking performance was evaluated by calculating the area under the error curve and by cross-correlation between the exerted and target paths. MU potential trains were then extracted and divided into ascending and descending segments for further analysis.

### Measurement of knee extensor strength

Participants were seated in a custom-built chair, which was adjusted to ensure approximately 90 degrees of flexion at the hips and knees. The lower leg was secured to a force dynamometer using a non-compliant strap (designed with a calibrated strain gauge, RS125 Components Ltd, Corby, UK) positioned above the medial malleolus. To minimize movement of the hips and upper trunk during contractions, the hips and pelvis were stabilised using a seat belt. Following a standardized warm-up consisting of submaximal contractions, participants were instructed to contract “as hard and as fast as possible” to ensure maximal effort with the aid of real-time visual feedback and verbal encouragement, aiming for forceful and rapid contractions. During the trial, participants were not allowed to hold onto the sides of the chair and were instructed to cross their arms over their chest. It was further repeated two to three times interspersed with 60 seconds rest intervals. If there was a difference of less than 5% between the two last attempts, the highest value was accepted as the maximal voluntary contraction (MVC).

### Muscle force control assessment

Muscle force control was evaluated during steady submaximal isometric contractions at 25% of MVC. Participants completed four trials following a trapezoidal profile (5s ramp-up, 12s plateau, 5s ramp-down), and the third trial was selected for detailed analysis as it represented a steady-state contraction free from initial familiarisation effects and potential fatigue observed in later trails. Muscle force signal was recorded at 2000Hz via a CED Micro 1401-3 (Cambridge Electronic Design, Cambridge, UK), interfaced with a desktop computer, with data exported for furtherr analysis in MATLAB (R2024b; The MathWorks, Natick, MA, USA). Force steadiness was quantified using the coefficient of variation (CoV), calculated as the standard deviation of the force during the steady plateau phase divided by its mean, multiplied by 100. The measure reflects the relative magnitude of force fluctuations.

To assess the ability to modulate force output in response to dynamic demands, participants performed a sinusoidal force tracking task using OTBioLab software (OT Bioelettronica, Turin, Italy). Prior to the test, a familiarisation trial was completed to ensure task understanding and reduce learning effects. The contraction included a 10s linear ramp-up phase, followed by eight sinusoidal oscillations at a fixed amplitude of 25 ± 4% MVC over 30s, and concluded with a 10s ramp-down phase. The recorded sinusoidal contractions were exported and analysed using MATLAB software (R2024b; The MathWorks, Natick, MA, USA). Force signals were low-pass filtered using a fourth-order Butterworth filter with a cutoff frequency of 20Hz prior to analysis. To quantify force tracking accuracy, the absolute error between the exerted path and the target path was calculated over the entire sinusoidal phase. The area under this error curve (AUC) was then calculated and normalised to the total duration of the contraction, yielding a normalised AUC (NormAUC, in N). Larger NormAUC values indicate greater tracking error, thus reflecting lower force tracking accuracy. The temporal coupling between target and exerted force paths during sinusoidal contractions was quantified using cross-correlation analysis, with the maximum Pearson correlation coefficient (r) used as an index of synchrony.

### High-density Surface Electromyography

High-density surface electromyography (HDsEMG) signals were recorded from the vastus lateralis muscle during the 25% of MVC trapezoidal contraction and sinusoidal force tracking task. A semi-disposable HDsEMG array (64 electrodes, 13×5, 8mm, I.E.D., GR08MM1305, OT Bioelettronica, Inc., Turin, Italy) was positioned over the muscle belly of the vastus lateralis with approximate orientation of the muscle fascicles (proximal to distal). Prior to placement, the skin was prepared by shaving, lightly abrading, and cleansing with 70% ethanol. The electrodes were affixed to the skin using flexible tape, and disposable bi-adhesive foam layers (SpesMedica, Bettipaglia, Italy) were used to secure the adhesive grids to the muscle surface. The skin electrode contact was facilitated by filling the cavities of the adhesive layers with conductive paste (AC Cream, SpesMedica). A strap ground electrode (WS2, OTBioelettronica, Turin, Italy) moistened with water was placed around the ankle of the tested leg. The signals were acquired in a monopolar configuration, amplified (x256), and filtered within the range of 10-500 Hz. The signals were then digitally converted at a sampling rate of 2000 Hz using a 16-bit wireless amplifier (Sessantaquattro, OTBioelettronica, Turin, Italy) and stored for further offline processing.

The recorded HDsEMG signals were converted into MATLAB format, and monopolar signals were further band-pass filtered (20 to 500 Hz) before decomposition into motor unit potential trains (MUPTs) using a convolutive kernel compensation (CKC) algorithm (Holobar & Zazula, 2007). All decomposed MUPTs were manually reviewed by a trained investigator to ensure accuracy of discharge patterns. Only MUs with a pulse-to-noise ratio equal to or greater than 28 dB were retained for further analysis. For sinusoidal contractions, only the oscillatory section of the signal was analysed, excluding the initial ramp-up and final ramp-down. Each full oscillation was further segmented into ascending and descending phases for phase-specific analysis.

Motor unit firing rate (MUFR) was calculated as the mean instantaneous discharge rate during the plateau phase of trapezoidal contractions and the oscillation cycle of sinusoidal contractions, with phase-specific analysis performed separately for ascending and descending segments. Motor unit firing rate variability (FRV) was quantified as the coefficient of variation (CoV) of the inter-spike intervals (ISI) during the plateau. MUFR at recruitment (FRrec) and derecruitment (FRderec) phases was defined as the mean instantaneous discharge rate across the first five discharges. Recruitment and derecruitment thresholds (RecThrsh and DerecThrsh) were defined as the force level corresponding to the first and final detected firing of each motor unit, respectively, and expressed as a percentage of the individual’s maximal voluntary contraction (%MVC).

### Statistical Analysis

All statistical analysis was conducted using RStudio. Multilevel mixed-effects linear regression models were used through the package lme4 (Version 1.1-27.1) (Bates *et al*., 2015) with each individual being regarded as an independent cluster.

To explore the bilateral differences in young and older individuals, Leg and AgeGroup were included as categorical variables, and their interactions (Leg × AgeGroup) were explored in the models. Maximal voluntary contraction (MVC) was included as a covariate in all models to control for inter-individual differences in strength. For the sinusoidal contractions, the waveform was segmented into ascending and descending phases to explore potential phase-dependent modulations in MU behaviour. Accordingly, SegType was included as a fixed factor in the overall model, and additional models were run separately for each phase to further examine within-task differences. Because multiple sinusoidal segments were obtained from each participant, models included a random intercept for both Subject and Segment (nested within Subject) to account for repeated measures. Normality assumptions were assessed visually using Q–Q plots of model residuals.

Where the interaction term was statistically significant, simple effects analyses were conducted using estimated marginal means (EMMs) (Lenth, 2016) to examine the effect of Leg within each AgeGroup (and vice versa). If the interaction was not significant, main effects of AgeGroup and Leg were interpreted directly based on the model estimates.

To assess model fit and estimate the proportion of variance explained, the marginal R^2^ (R^2^_m_) and conditional R^2^ (R^2^c) are included in each table of model outputs. The R^2^_m_ represents the proportion of variance explained by the fixed effects, and R^2^c represents the variance explained by both fixed effects and random effects (Nakagawa & Schielzeth, 2013). Cohen’s d effect size was calculated from EMMs derived from the linear mixed-effects models, accounting for the nested data structure. Commonly used interpretations separate effects sizes intro three categories: small (d = 0.2), medium (d = 0.5) and large (d = 0.8). Statistical significance was accepted when p<0.05.

## Results

A total of 38 healthy participants were included in the study, consisting of 21 younger adults (15 males, 6 females; age range: 20-35 years) and 17 older adults (12 males, 5 females; age range: 67-83 years). MU data meeting the analysis criteria were successfully obtained from 18 younger and 17 older participants. Across all participants, a total of 940 MUs were identified during trapezoidal contractions and 584 during sinusoidal contractions (Table 1).

**Table 1.** Participant characteristics.

| Measurements | Younger |  | Older |  |
| --- | --- | --- | --- | --- |
|  | Dominant | Non-Dominant | Dominant | Non-Dominant |
| Age (years) | 22 ± 4 |  | 74 ± 5 |  |
| Body mass index (kg/m²) | 24.27 ± 3.62 |  | 25.57 ± 2.95 |  |
| *Muscle force control* |  |  |  |  |
| MVC (N) | 510.64 ± 159.32 | 495.08 ± 178.57 | 281.68 ± 94.47 | 276.81 ± 89.88 |
| CoV-Force | 2.50 ± 0.92 | 2.62 ± 1.23 | 2.56 ± 0.78 | 2.18 ± 0.87 |
| Sin NormAUC | 0.77 ± 0.18 | 0.85 ± 0.22 | 1.14 ± 0.41 | 1.13 ± 0.39 |
| Sin CC | 0.94 ± 0.03 | 0.93 ± 0.03 | 0.89 ± 0.06 | 0.88 ± 0.06 |
| *Motor unit properties - Trapezoidal Contraction* |  |  |  |  |
| N (MUs) | 245 | 180 | 244 | 271 |
| MUFR (Hz) | 8.67 ± 1.24 | 8.68 ± 1.48 | 7.57 ± 1.10 | 7.64 ± 1.10 |
| MUFR Variability (%) | 12.32 ± 3.88 | 15.70 ± 9.91 | 11.40 ± 3.01 | 12.77 ± 3.84 |
| RecThrsh (%MVC) | 12.04 ± 2.22 | 12.39 ± 2.57 | 11.16 ± 2.35 | 11.37 ± 1.60 |
| DerecThrsh (%MVC) | 13.50 ± 2.73 | 12.87 ± 3.02 | 12.57 ± 4.27 | 12.58 ± 3.71 |
| *Motor unit properties - Sinusoidal Contraction* |  |  |  |  |
| N (MUs) | 169 | 131 | 148 | 134 |
| MUFR (Hz) | 8.98 ± 1.37 | 8.93 ± 1.20 | 7.66 ± 1.25 | 7.76 ± 1.17 |
| MUFR Variability (%) | 21.57 ± 5.02 | 18.39 ± 4.23 | 16.40 ± 3.57 | 17.95 ± 4.10 |

*Data are reported as mean ± standard deviation. Abbreviations: MVC, maximal* *voluntary isometric contraction; CoV, coefficient of variation; Sin NormAUC,* *normalised area under sinusoidal curve; Sin CC, cross-correlation coefficients* *between sinusoidal curves; MUFR, motor unit firing rate; RecThrsh, recruitment* *threshold; DerecThrsh, derecruitment threshold.*

### Muscle Force Control

For MVC, there was a significant main effect of AgeGroup (β = -1.29, Δ = -228.97 N, 95% CI [-1.81 to -0.78], p < 0.001), with older adults generating lower maximal force compared to younger adults. There was no significant Leg effect (β = -0.09, Δ = - 15.56 N, 95% CI [-0.27 to 0.10], p = 0.344) nor a significant Leg × AgeGroup interaction (p = 0.663) (Table 2, Figure 2A). In the analysis of force steadiness (CoV-Force) during the trapezoidal contractions, no significant Leg × AgeGroup interaction was observed (p=0.097). There was also no significant main effect of AgeGroup (β = 0.06, Δ = +0.06, 95% CI [-0.59 to 0.71], p = 0.851), nor significant main effect of Leg (β = 0.13, Δ = +0.12, 95% CI [-0.29 to 0.54], p = 0.549) (Table 2, Figure 2B).

**Figure 2.**
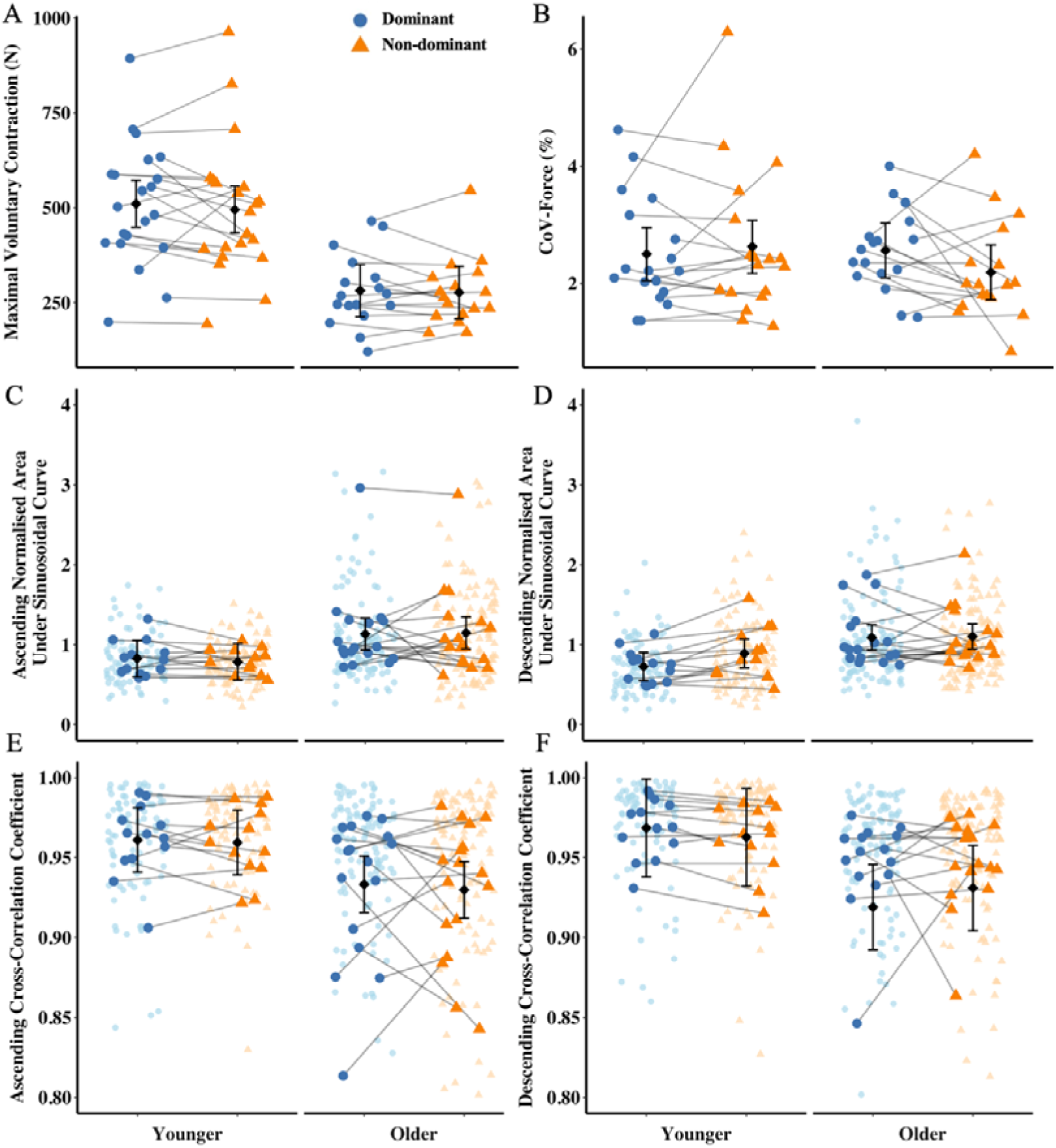
Measures of muscle force control in younger and older adults. Panels show maximal voluntary contraction (MVC) (A), coefficient of variance of force (CoV-Force) (B) during trapezoidal contractions, normalised area under the sinusoidal force curve (C-D) and cross-correlation coefficient (E-F) between target and exerted force during ascending and descending segments of sinusoidal contractions. Each data point represents each leg of an individual participant, with dominant and non-dominant legs indicated by purple circles and blue triangles, respectively. Individual motor unit data across segments are plotted as jittered points in panels C-F. Mean values for each leg per participant are connected by lines to visualise within subject differences across legs. Black dots with error bars represent estimated marginal means (EMMs) ± standard errors from the statistical model.

**Table 2.** Summary of mixed-effects linear model results for muscle force control measures.

| Measures | Leg (NonDom v Dom) | AgeGroup<br>(Young v Old) | Leg × AgeGroup | Model Fit<br>($R^2m/R^2c$) |
| --- | --- | --- | --- | --- |
| MVC (N) | $\leftrightarrow -15.56$ ($\beta=-0.09$,<br>$p=0.344$) | $\downarrow -228.97$ ($\beta=-1.29$, $p<0.001$) | $p=0.663$ | 0.390 / 0.913 |
| CoV-Force (%) | $\leftrightarrow 0.13$ ($\beta=0.12$,<br>$p=0.549$) | $\leftrightarrow -0.06$ ($\beta=-0.06$, $p=0.851$) | $p=0.097$ | 0.030 / 0.595 |
| Sin NormAUC | $\leftrightarrow 0.06$ ($\beta=0.11$,<br>$p=0.155$) | $\uparrow 0.33$ ($\beta=0.63$, $p=0.008$) | $p=0.409$ | 0.085 / 0.449 |
| Sin CC | $\leftrightarrow -0.00$ ($\beta=-0.03$,<br>$p=0.743$) | $\downarrow -0.04$ ($\beta=-0.35$, $p=0.007$) | $p=0.592$ | 0.024 / 0.082 |

**Note.** Data presented as fixed effect estimates ( $\beta$ ) with corresponding $p$ values. Arrows indicate the direction of the effect: ↑ increase, ↓ decrease, ↔ no difference. The final column shows model fit statistics. Abbreviations: MVC, maximal voluntary contraction; CoV-Force, coefficient of variation of force; Sin NormAUC, normalised area under the sinusoidal curve; Sin CC, correlation coefficient during sinusoidal tracking; $R^2m$ , marginal $R^2$ representing variance explained by fixed effects only; $R^2c$ , conditional $R^2$ representing variance explained by the full model (fixed + random effects). \*Statistically significant ( $p < 0.05$ ).

In the segmented sinusoidal force-tracking task, no significant Leg × AgeGroup interaction was observed for either normalised area under the curve (NormAUC) (p=0.409) or cross-correlation coefficients (CC) (p=0.592). There were also no significant main effects of Leg (NormAUC: β = 0.11, p = 0.155; CC: β = -0.03, p = 0.743) or Segment Type (NormAUC: β = -0.05, p = 0.374; CC: β = -0.01, p = 0.854) on tracking accuracy. However, significant main effects of AgeGroup were found for both outcome measures, with older adults showing higher normalised AUC values (β = 0.63, Δ = +0.33, 95% CI [0.09 to 0.58], p = 0.008) and lower cross-correlation coefficients (β = -0.35, Δ = -0.04, 95% CI [-0.07 to -0.01], p = 0.007) (Table 2, Figure C-F).

### Motor Unit Properties During 25% MVC Trapezoidal Contractions

For MUFR, no significant Leg × AgeGroup interaction was observed (p = 0.779). A significant main effect of AgeGroup was found, with older adults exhibiting lower MUFR compared to younger adults (β = -0.64, Δ = -1.18 Hz, 95% CI [-1.03 to -0.26], p = 0.001). No significant main effect of Leg was detected (β = 0.01, Δ = 0.01 Hz, 95% CI [-0.12 to 0.14], p = 0.908) (Table 3, Figure 3A). For MUFR variability, there was no significant Leg × AgeGroup interaction (p = 0.102). A significant main effect of Leg was found (β = 0.28, Δ = 2.08%, 95% CI [0.13 to 0.43], p < 0.001), indicating greater variability in the non-dominant legs in both groups, but no significant main effect of AgeGroup (β = -0.01, Δ = -0.08%, 95% CI [-0.32 to 0.30], p = 0.948) (Table 3, Figure 3B). For recruitment and derecruitment thresholds, neither Leg nor AgeGroup showed significant main effects, and there was no significant interaction (all p > 0.20) (Table 3, Figure 3C-D).

**Figure 3.**
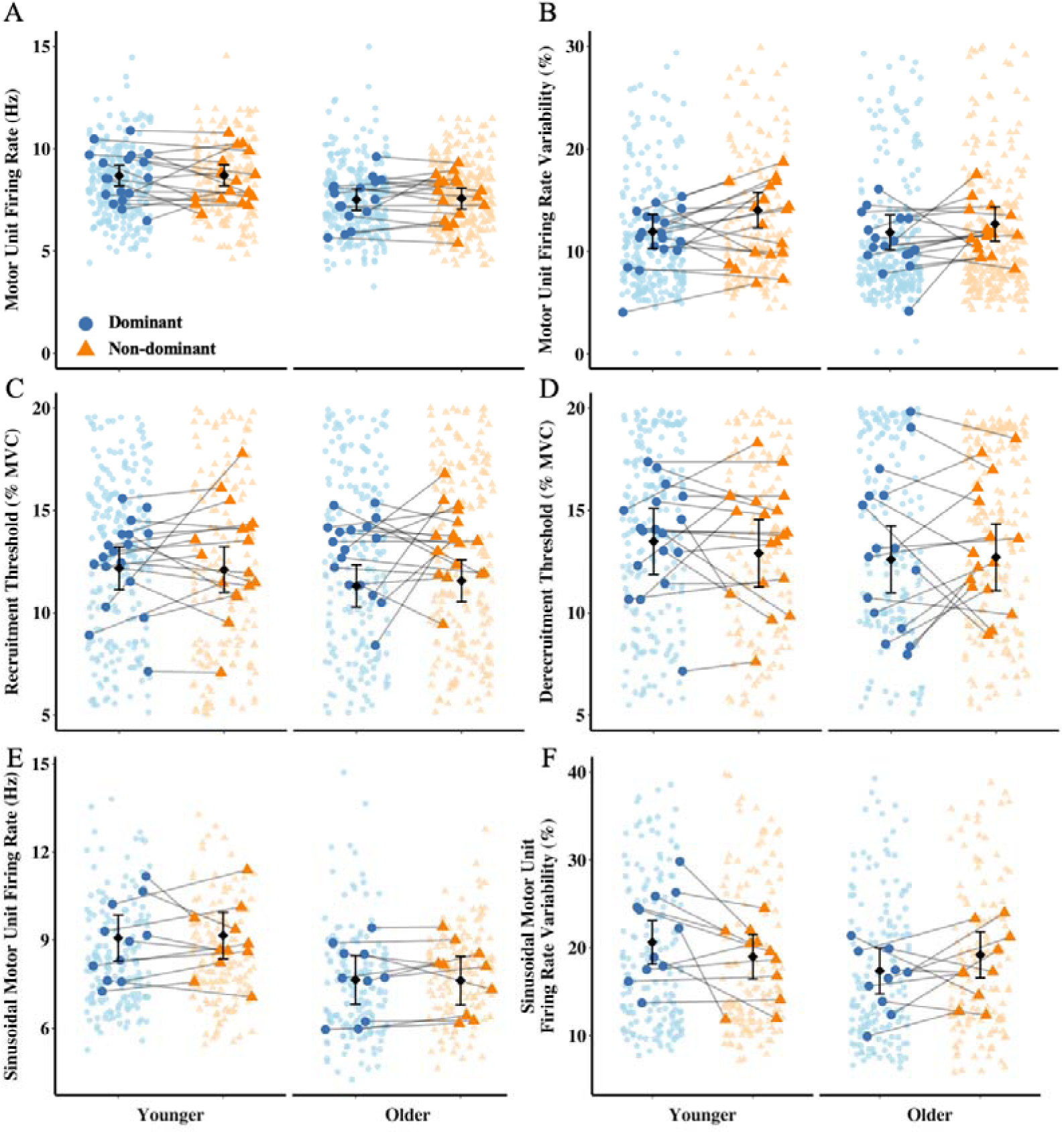
Motor unit firing properties in younger and older adults during trapezoidal and sinusoidal contractions. Panels A and B show motor unit firing rate (MUFR) and MUFR variability during 25% maximal voluntary trapezoidal contractions. Panel C and D illustrate recruitment and derecruitment thresholds, respectively. Panel E and F present MUFR and MUFR variability during sinusoidal force-tracking tasks. Data are presented separately by AgeGroup (Younger vs older adults) and Leg dominance (Dominant vs non-dominant). Purple circles are representing dominant legs and blue triangles represent non-dominant legs. Individual motor unit data are plotted as jittered points. Mean values for each leg per participant are connected by lines to visualise within subject differences across legs. Black dots with error bars represent estimated marginal means (EMMs) ± standard errors from the statistical model.

**Table 3.** Summary of multilevel linear regression model results for motor unit firing properties.

| Contraction Types | Measures | Leg (NonDom v Dom) | AgeGroup (Old v Young) | AgeGroup × Leg | Model Fit (R<sup>2</sup>m/R<sup>2</sup>c) |
| --- | --- | --- | --- | --- | --- |
| **Trapezoidal Contraction** | MUFR (Hz) | ↔ 0.01 ($\beta=0.01$, $p=0.908$) | ↓ -1.18 ($\beta=-0.64$, $p=0.001$) | $p=0.779$ | 0.360 / 0.628 |
| | MUFR Variability (%) | ↑ 2.08 ($\beta=0.28$, $p<0.001$) | ↔ -0.08 ($\beta=-0.01$, $p=0.948$) | $p=0.102$ | 0.290 / 0.450 |
| | RecThrsh (%MVC) | ↔ -0.07 ($\beta=-0.01$, $p=0.899$) | ↔ -0.86 ($\beta=-0.16$, $p=0.237$) | $p=0.651$ | 0.004 / 0.083 |
| | DerecThrsh (%MVC) | ↔ -0.58 ($\beta=-0.10$, $p=0.215$) | ↔ -0.88 ($\beta=-0.16$, $p=0.438$) | $p=0.267$ | 0.004 / 0.313 |
| **Sinusoidal Contraction** | MUFR (Hz) | ↑ 0.26 ($\beta=0.13$, $p<0.001$) | ↓ -1.10 ($\beta=-0.56$, $p=0.061$) | $p=0.057$ | 0.142 / 0.541 |
| | MUFR Variability (%) | ↓ -0.06 ($\beta=-0.20$, $p<0.001$) | ↓ -0.10 ($\beta=-0.34$, $p=0.002$) | $p<0.001$ | 0.085 / 0.167 |

***Note.** Data are presented as fixed effect estimates ( $\beta$ ) with corresponding $p$ values.* *Arrows indicate the direction of the effect: ↑ increase, ↓ decrease, ↔ no meaningful* *difference. The final column shows model fit statistics. Abbreviations: MUFR, motor* *unit firing rate; MUFR variability, coefficient of variation of motor unit firing rate;* *RecThrsh, recruitment threshold; DerecThrsh, derecruitment threshold; $R^2$ ,* *marginal $R^2$ (variance explained by fixed effects only); $R^2c$ , conditional $R^2$ (variance* *explained by the full model including both fixed and random effects). Statistically* *significant ( $p < 0.05$ ).*

### Motor Unit Properties During Sinusoidal Contractions

No significant Leg × AgeGroup interaction in MUFR was observed (p = 0.057). There was a significant main effect of Leg, indicating higher MUFR in the non-dominant limb (β = 0.13, Δ = 0.26 Hz, 95% CI [0.08 to 0.19], p < 0.001), and a nonsignificant trend toward lower MUFR in older adults (β = -0.56, Δ = -1.10 Hz, 95% CI [-1.15 to 0.03], p = 0.061) (Table 3, Figure 3E). For MUFR variability, a significant Leg × AgeGroup interaction was observed (p < 0.001). Main effects were significant for both Leg (β = -0.20, Δ = -0.06%, 95% CI [-0.27 to -0.14], p < 0.001) and AgeGroup (β = -0.34, Δ = -0.10%, 95% CI [-0.56 to -0.12], p = 0.002). Post-hoc analysis revealed that within-group comparisons, younger adults had higher MUFR variability in their dominant leg (p < 0.001, d = 0.26), while older adults showed a lower variability in their dominant legs (p = 0.009, d = -0.09) (Table 3, Figure 3F).

In addition to the main effects of Leg and AgeGroup, significant main effects of segment type were observed in MUFR and MUFR variability during the sinusoidal contractions. Specifically, MUFR was higher during ascending phases (β = 0.27, 95% CI [0.42 to 0.12], p < 0.001), and MUFR variability was markedly greater during descending segments (β = 0.14, 95% CI [0.11 to 0.17], p < 0.001). These consistent effects highlight the importance of analysing ascending and descending phases separately when evaluating age- and leg-related differences in MU behaviour.

When phases were explored separately, the ascending phase of sinusoidal contractions showed no significant Leg × AgeGroup interaction for MUFR (p = 0.096). There was a significant main effect of Leg (β = 0.22, Δ = 0.43 Hz, 95% CI [0.15 to 0.29], p < 0.001), with higher MUFR in non-dominant legs, whereas there was no significant main effect of AgeGroup (β = -0.38, Δ = -0.73 Hz, 95% CI [-1.04 to 0.29], p = 0.272) (Table 4, Figure 4A). For MUFR variability, a significant Leg × AgeGroup interaction was found (p < 0.001), along with significant main effects of Leg (β = -0.22, Δ = -0.02%, 95% CI [-0.31 to -0.13], p < 0.001) and AgeGroup (β = -0.62, Δ = -0.07%, 95% CI [-1.04 to -0.20], p = 0.004). Post-hoc analysis showed that MUFR variability during ascending phases was higher in the dominant legs compared to the non-dominant legs in younger adults (Δ = 0.02%, p < 0.001, d = 0.24), while older adults showed a reversed but non-significant pattern (Δ = -0.01%, p = 0.099, d = -0.08). Age-related differences revealed greater MUFR variability in younger adults than older adults, with more pronounced differences in the dominant legs (Δ = 0.07%, p = 0.004, d = 0.66) than in the non-dominant legs (Δ = 0.03%, p = 0.13, d = 0.35) (Table 4, Figure 4C).

**Figure 4.**
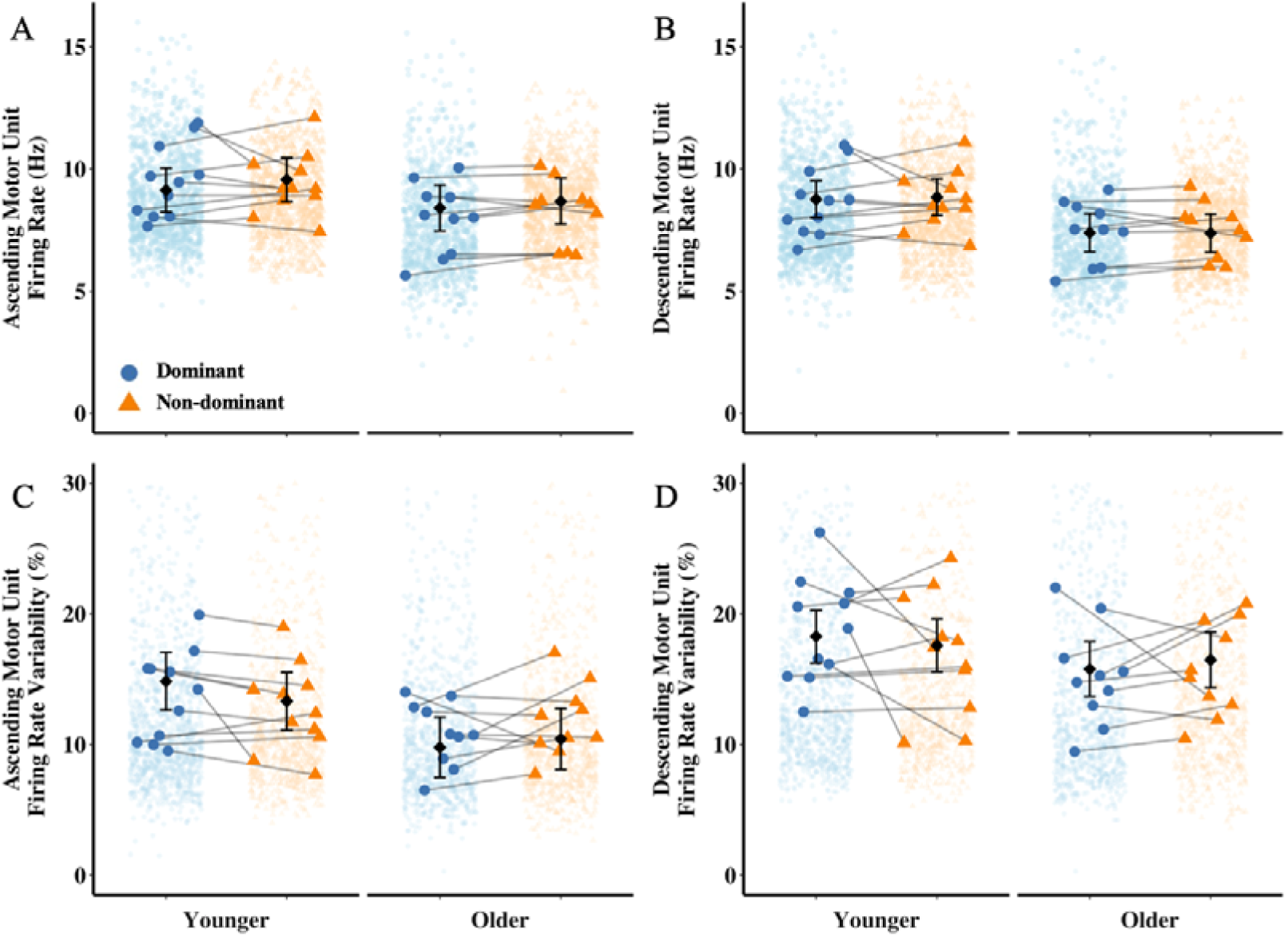
Motor unit firing properties during ascending and descending phases of sinusoidal contractions in younger and older adults. Panels A and C show motor unit firing rate (MUFR) and MUFR variability during the ascending phase of a sinusoidal contraction. Panels B and D display the corresponding measures during the descending phase. Data are presented separately by AgeGroup (Younger vs older adults) and Leg dominance (Dominant vs non-dominant). Purple circles are representing dominant legs and blue triangles represent non-dominant legs. Individual motor unit data are plotted as jittered points. Mean values for each leg per participant are connected by lines to visualise within subject differences across legs. Black dots with error bars represent estimated marginal means (EMMs) ± standard errors from the statistical model.

**Table 4.**
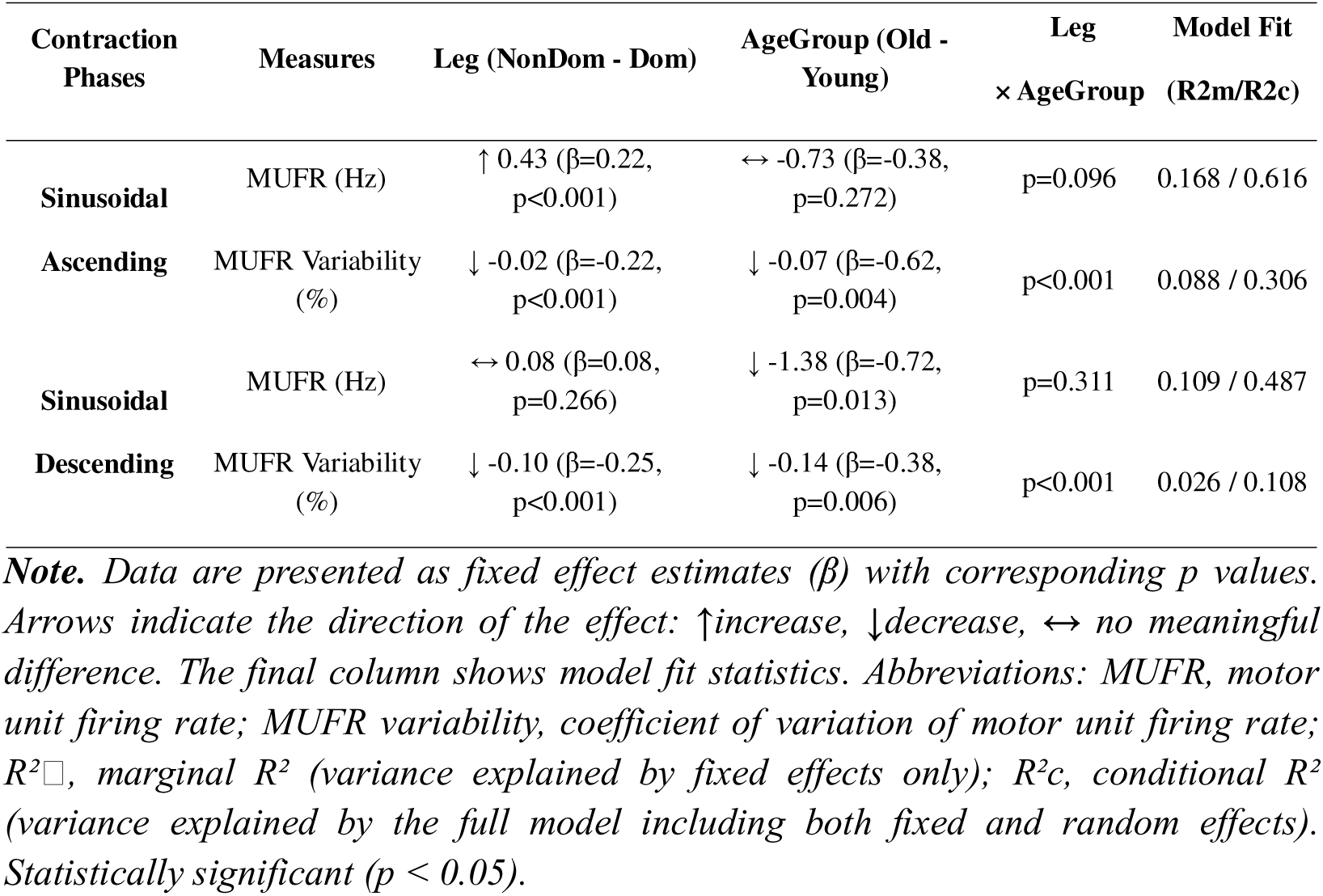
Summary of multilevel linear regression model results for motor unit firing properties during ascending and descending phases of sinusoidal contractions.

During the descending phase of sinusoidal contractions, no significant Leg × AgeGroup interaction was found in MUFR (p = 0.311) nor a main effect of Leg (β = 0.08, Δ = 0.08 Hz, 95% CI [-0.03 to 0.12], p = 0.266). However, there was a significant main effect of AgeGroup (β = -0.72, Δ = -1.38 Hz, 95% CI [-1.29 to -0.15], p = 0.013), with lower MUFR in older adults across both legs (Table 4, Figure 4B). For MUFR variability, a significant Leg × AgeGroup interaction was found (p < 0.001), along with main effects of Leg (β = -0.25, Δ = -0.10%, 95% CI [-0.34 to -0.16], p < 0.001) and AgeGroup (β = -0.38, Δ = -0.14%, 95% CI [-0.65 to -0.11], p = 0.006). Post-hoc analyses showed that MUFR variability was greater in the dominant leg compared to the non-dominant leg in younger adults (Δ = 0.10%, p < 0.001, d = 0.26), whereas older adults exhibited the opposite pattern (Δ = –0.04%, p = 0.025, d = –0.10). Age-related differences were more pronounced in the dominant leg (Δ = 0.14%, p = 0.006, d = 0.40) than in the non-dominant leg (Δ = 0.01%, p = 0.84, d = 0.03) (Table 4, Figure 4D).

## Discussion

The present study investigated bilateral differences in muscle force control and MU firing behaviours in young and older adults across constant and variable force tracking tasks. Our key findings revealed clear age-related impairments in neuromuscular function, including reduced maximal voluntary force and reduced accuracy during sinusoidal force tracking tasks. No significant leg asymmetries were observed in either age group, suggesting age-related declines in neuromuscular function are bilateral and relatively symmetric. Notably, contraction type and age influenced MU firing patterns, with variability differing across phases. These findings suggest that leg-related asymmetries in MU firing behaviour are exacerbated with ageing, particularly during complex dynamic force modulation.

Our study observed no significant bilateral differences in knee extensor strength in either age group. This aligns with previous findings in other lower limb muscles, such as the tibialis anterior (Petrovic *et al*., 2022) and likely reflects routine bilateral engagement of the lower limbs. Although numerous studies have reported reduced force steadiness in older adults compared to younger counterparts (Oomen & van Dieën, 2017; Pethick *et al*., 2022), we found no significant age-related differences in force steadiness, possibly due to the relatively high health and functional status of our older participants. The observed bilateral symmetry in simple tasks suggests that isometric force control relies on generalized mechanisms rather than limb-specific adaptations, or, that limb-specific adaptations in lower limbs do not occur. However, this symmetry did not extend to more difficult sinusoidal force tracking tasks, where older adults exhibited significantly reduced performance. This dissociation highlights the task-specific nature of age-related declines, as the demands of variable force tracking strongly rely on mechanosensitive afferents and cognitive-motor coupling, known to decrease with age (Störmer *et al*., 2013; McIntyre *et al*., 2021). Thus, the impaired tracking performance observed in older adults may partially reflect age-related central limitations, beyond peripheral neuromuscular deficits. While these force-related findings suggest age-related differences in control strategies, they provide limited insight into the underlying neural mechanisms.

MUFR exhibited task- and age-specific modulation, with more pronounced effects during variable force contractions. In constant force contractions, MUFR did not differ between legs in either age group, consistent with prior findings in lower limbs (e.g., tibialis anterior) (Petrovic *et al*., 2022), whereas limb-specific differences have been reported in the upper limbs (e.g., biceps brachii) (Lecce *et al*., 2025). However, older adults exhibited significantly lower MUFR, which is a widely reported finding in both upper and lower limbs (Orssatto *et al*., 2022). These results suggest that under steady contractions, MUFR is primarily influenced by age, rather than leg dominance. In contrast, sinusoidal (variable force) contractions revealed more complex modulation patterns. To better understand these task-dependent modulations, sinusoidal contractions were further segmented into ascending (concentric) and descending (eccentric) phases. MUFR was consistently higher during the ascending phase, reflecting the increased neural drive required for force generation. These data align with prior findings demonstrating that MUFRs tend to be greater during concentric than eccentric contractions, even within the same MUs, especially during low-force dynamic task (Del Valle & Thomas, 2005; Duchateau & Baudry, 2013; Basile *et al*., 2025). The greater MUFR during the ascending phase may be attributable to enhanced corticospinal excitability and greater neuromuscular recruitment demands associated with concentric movement, potentially also reflecting reduced inhibitory control compared to eccentric contractions (Gueugneau *et al*., 2023).

Importantly, when examining sinusoidal contractions in detail, a distinct modulation pattern related to leg dominance was observed, which was not present during constant force tasks. Specifically, MUFR during the ascending phase was significantly higher in the non-dominant legs in both age groups. This finding partially aligns with some prior work but contrasts with others. While several studies have reported higher MUFR on the dominant side across different contraction intensities (Lecce *et al*., 2025), others have found no difference (Petrovic *et al*., 2022) or even non-dominant advantages (Adam *et al*., 1998). These discrepancies may arise from differences in muscle selection, contraction type and the inherently dynamic nature of sinusoidal contractions, which engaged greater neuromuscular demands than constant plateaus. The present findings support the presence of lateralised neuromuscular control strategies that influence force generation more than force reduction. The ascending phase, characterised by increasing force output, relies heavily on corticospinal drive (Clos *et al*., 2022). While the dominant limbs are associated with stronger neural drive (Lecce *et al*., 2025) and more effective proprioceptive feedback (Strong *et al*., 2023), the higher MUFR observed in the non-dominant leg may indicate a compensatory strategy, possibly involving changes in inhibition control or motoneuron excitability. These results collectively suggest that leg dominance interacts with task dynamics to shape neuromuscular recruitment strategies, particularly during phases of increasing force.

When examining these phase-specific effects across age groups, during the descending (eccentric) phase, MUFR was lower in older adults. Previous studies have highlighted that eccentric contractions rely more heavily on cortical involvement and require greater sensorimotor integration than concentric contractions (Duchateau & Baudry, 2013). However, ageing is associated with reduced corticospinal excitability (Oliviero *et al*., 2006), particularly during eccentric tasks (Duclay *et al*., 2014; Clos *et al*., 2022; Ruas *et al*., 2025), which likely contributes to impaired discharge modulation during muscle lengthening contractions. Additionally, the reduced MUFR during the descending phase could reflect not only impaired excitatory drive but also enhanced inhibitory processes within the motor system, Specifically, older adults may exhibit increased presynaptic inhibition at the spinal level (Baudry & Duchateau, 2012; Filho *et al*., 2021) or reduced intracortical facilitation during eccentric movements (Adam *et al*., 1998), both of which could further limit the ability to modulate MU firing pattern. This suggests that the age-related decline in neural drive is more pronounced during phases of force reduction than force generation. This finding has important considerations for activities of daily living which rely on eccentric contraction, such as stair descent.

While MUFR serves as a fundamental parameter for investigating neuromuscular recruitment strategies, its variability offers additional insights into the neuromuscular stability and adaptability, influenced by task difficulty, leg dominance and age-related changes. In the present study, during constant load contractions, both age groups showed greater MUFR variability in the non-dominant legs. This aligns with earlier findings suggesting greater MUFR fluctuations on the non-dominant side (Adam *et al*., 1998), potentially reflecting a stronger reliance on afferent sensory feedback and spinal-level regulation in that limb. During sinusoidal contractions, which require continuous force modulation and more dynamic engagement of higher-order motor control systems, MUFR variability was markedly greater across all participants. This task-dependent increase in MUFR variability highlights the increased demands placed on neuromuscular coordination and sensorimotor integration throughout the contraction.

MUFR variability is generally higher with higher MUFR in young and older VL (Guo et al., 2022, 2025), but the current data show this relationship dissociates when comparing ascending and descending segments. Despite lower MUFR, the descending (eccentric) phase exhibited significantly higher MUFR variability than the ascending (concentric) phase. This suggests that MUFR variability reflects distinct phase-specific neural control strategies, not just firing rate.The increased variability during eccentric contraction may reflect stronger inhibitory regulation, such as presynaptic inhibition or cortical inhibition, which are known to be upregulated during lengthening eccentric contractions (Doguet *et al*., 2017; Papitsa *et al*., 2022). However, age-related impairments in muscle relaxation (Hunter *et al*., 2016; Jeon *et al*., 2023) may further limit motor output stability in older adults during this phase.

Age-related differences in MUFR variability became more pronounced during sinusoidal contractions. In younger adults, MUFR variability was higher in the dominant leg. This greater MUFR variability may be indicative of a more variable use of corticospinal resources by the dominant legs, such as that proposed following the finding of fewer common synaptic inputs in dominant hands (Maillet *et al*., 2022), thereby enabling task-specific adjustments to motor output. Such flexibility may be particularly advantageous in tasks involving variable forces, as it facilitates rapid transition in force. However, this was not apparent in the older adults, where MUFR variability was more similar across legs, suggesting a shift towards more generalised control strategies. This phenomenon may be attributable to age-related corticospinal degradation or diminished inter-hemispheric communication (Delvenne & Malloy, 2025). Notably, older adults demonstrated reduced bilateral modulation across tasks, with smaller inter-limb differences in MUFR variability during variable force contractions. Overall, these findings suggest the progressive reduction in neuromuscular adaptability with aging, which may reflect a reduced ability to implement differentiated neural strategies across task-specific motor demands.

## Strengths and Limitations

This study investigated the effects of leg dominance on neuromuscular function across young and older people, including both static trapezoidal contractions and dynamic sinusoidal contractions. However, this study still has some limitations. (1) Both males and females were included in the study, but the sample size for female MU data was relatively small. (2) MUs sampled in the current study were only from mid-level contraction intensity, so revealed limited information on higher threshold MUs. (3) Knee extensor movements are controlled by multiple muscles, so investigating only the MUs of vastus lateralis MUs may limit the interpretation of neural control for the entire quadriceps muscle group.

## Conclusions

Our findings collectively indicate that the aging neuromuscular system exhibits heterogeneous adaptations rather than a uniform decline, with MU firing behaviours showing task- and leg-specific modulation. Despite the largely symmetrical use of lower limbs, subtle lateralised differences are evident, particularly during dynamic tasks requiring continuous modulation of neural drive. Notably, we also report distinct differences in MU behaviour across contraction types, with lower MUFR and higher variability during eccentric phases. Thus, instead of a uniform decline, ageing appears to reshape motor control through altered lateralised and phase-dependent strategies. These insights may help to inform the development of more targeted interventions to support functional independence in older adults.

## Acknowledgements

The authors thank all of the participants for their time and participation.

## Author Contributions

All authors contributed to the conception and design of the study. YG, AA, NNG, and EJ contributed to the data acquisition. YG and MP analysed the data and drafted the manuscript. BEP and PJA provided comments. All authors have approved the final version of the submitted manuscript for publication and are accountable for all aspects of the work. All persons designated as authors qualify for authorship, and all those who qualify for authorship are listed.

## Data availability statement

The datasets generated and analysed during the current study are available from the corresponding author upon reasonable request.

